# Bots for Software-Assisted Analysis of Image-Based Transcriptomics

**DOI:** 10.1101/172296

**Authors:** Marcelo Cicconet, Daniel R. Hochbaum, David Richmond, Bernardo L. Sabatini

**Affiliations:** Harvard Medical School, Boston, MA

## Abstract

We introduce software assistants – bots – for the task of analyzing image-based transcriptomic data. The key steps in this process are detecting nuclei, and counting associated puncta corresponding to labeled RNA. Our main release offers two algorithms for nuclei segmentation, and two for spot detection, to handle data of different complexities. For challenging nuclei segmentation cases, we enable the user to train a stacked Random Forest, which includes novel circularity features that leverage prior knowledge regarding nuclei shape for better instance segmentation. This machine learning model can be trained on a modern CPU-only computer, yet performs comparably with respect to a more hardware-demanding state-of-the-art deep learning approach, as demonstrated through experiments. While the primary motivation for the bots was image-based transcriptomics, we also demonstrate their applicability to the more general problem of scoring “spots” in nuclei.

## 1. Introduction

Image-based transcriptomics is a burgeoning field of research with applications from basic cell biology [4, 10] to systems neuroscience [25, 22]. *In situ* imaging methods for scoring^1^ fluorescently labeled RNA transcripts are attractive tools for studying biological tissues with complex and heterogeneous spatial environments that can signal and shape gene expression in single cells. However, scoring the number of fluorescently labeled RNA in each cell in a tissue remains a challenging bio-image analysis task.

Generally speaking, scoring fluorescent puncta in nuclei requires the combination of two independent image processing pipelines – one for nuclei segmentation, and one for puncta detection. Each pipeline presents its own inherent complexities: nuclei segmentation is a challenging case of instance segmentation [23], which requires both labeling pixels belonging to nuclei, as well as separating touching nuclei. Spot detection is also a challenging task under low signal-to-noise ratio conditions [1]. Currently, cell segmentation is most often achieved either by painstaking manual segmentation of fluorescent, DAPI-stained cell nuclei [22], or by narrowly defined and carefully tuned algorithms that are not readily generalizable or suitable for *in vivo* measurements [4], where cells can be clumped or clustered.

Here we present an open source pipeline for automated cell segmentation and puncta scoring. The pipeline relies on a stacked Random Forest model for challenging nuclei segmentation, and provides robust results, even with a small training set. We introduce novel “circularity features” to the stacked Random Forest model, to leverage the prior knowledge that nuclei are approximately spherical, such that the stacked Random Forest can learn to split touching nuclei. Our stacked Random Forest model with circularity features achieves a performance matching the popular U-Net model for nuclei segmentation, in a lightweight model that can be trained and run on CPU.

We make our solution available by way of *bots*, or software assistants, developed in Matlab. Bots use minimalistic GUI elements to guide the user through setting the various image-processing parameters. SpotsInNucleiBot (SNB), our main release, allows the user to set, load, and save parameters for segmenting nuclei, detecting puncta, and associating the two objects together (see Figure 1 for screenshots and sample workflow). It allows scoring one stack at a time, or its “headless” version can be used to score a folder of images as a traditional script on a PC or server. SNB offers two options for nuclei segmentation, and two for point-source detection, permitting a more adequate handling of data of different complexities. A basic algorithm can be used for simple, fast segmentation when the signal-to-noise ratio is high. For more challenging nuclei segmentation cases, SNB uses the multi-layer Random Forest model for pixel classification, followed by watershed segmentation of the resulting class probability maps. Both image annotation and model training can be done with an accompanying NucleiSegmentationBot (NSB).

**Figure 1.**
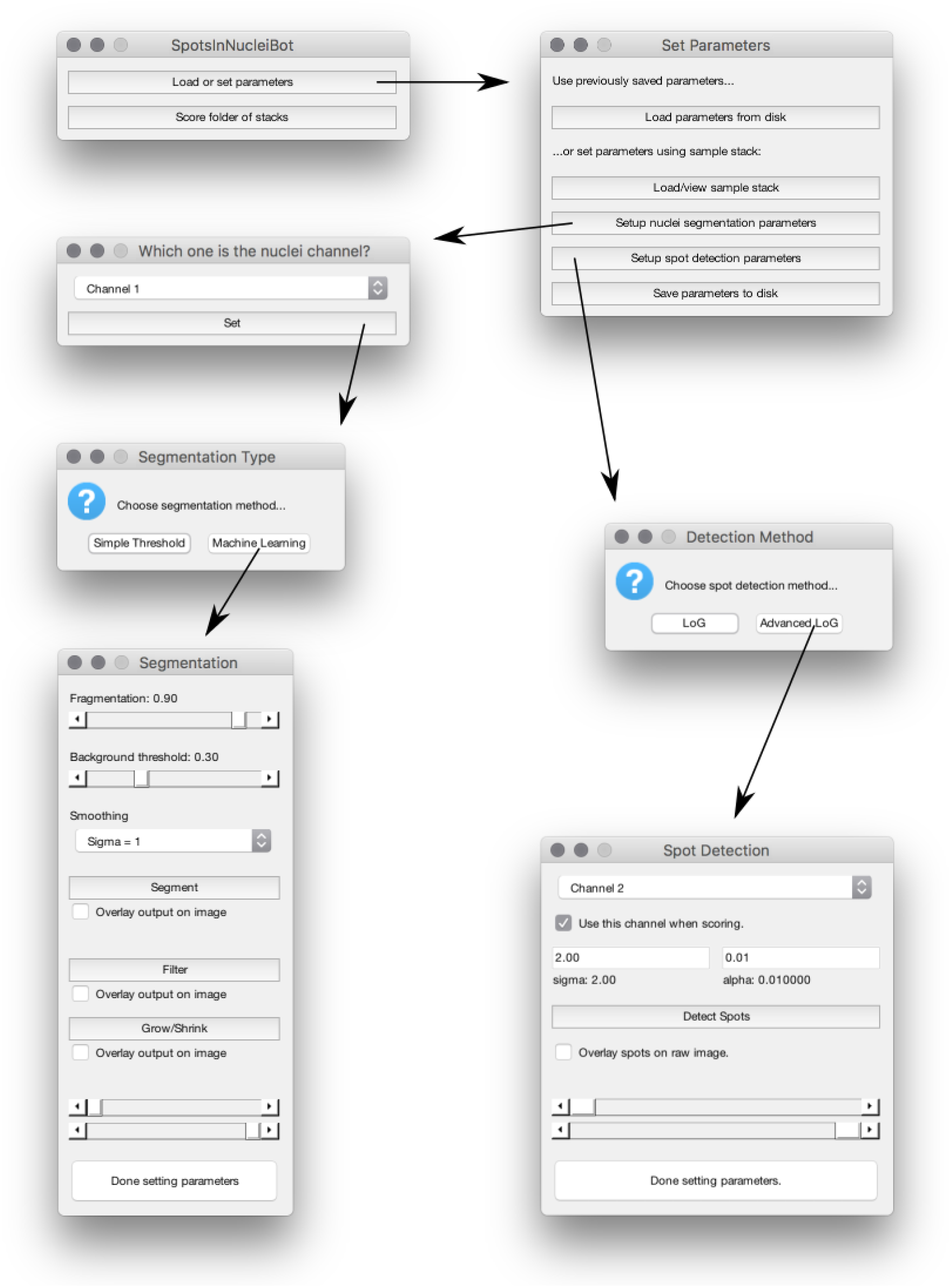
SpotsInNucleiBot sample workflow. The bot assists the user by asking questions via basic dialog windows. It also includes mini-apps (such as the Segmentation and Spot Detection windows above) that allow the user to experiment with parameters. Not shown is the Figure window, that displays the current image plane being analyzed, along with possible mask overlays (when that option is selected).

In addition to the SNB and NSB development, we are also releasing bots for estimating the parameters of individual puncta, building stacks from image planes, and visualizing stacks^2^. Source code for the Random Forest classifier using circularity features is also available independently^3^. We hope that the resource presented here will be useful for rapid hypothesis testing and validation for the increasingly diverse array of transcriptionally-defined cell types within complex tissues.

## 2. Related Work

There are many works addressing the task of nuclei segmentation in bio-image analysis. Methods based on classical computer vision include automatic thresholding [17, 19] combined with morphological operations, marker-controlled watershed [16, 27], active contours and level sets [9, 14], and model-based approaches such as the generalized hough transform [12, 3]. More recently, Deep Learning has dominated the field of semantic segmentation [15, 20], including instance segmentation [7, 2], whenever a large amount of annotated training data is available. We provide two methods for nuclei segmentation with our bots: thresholding-based segmentation for simple cases (e.g., well separated nuclei), as well as a Machine Learning based approach for more challenging cases frequently occurring in complex tissues. For the Machine Learning method, we chose to use a stacked Random Forest model as the classification back-end, rather than a Deep Learning model such as U-Net [20], because we wanted to develop a tool that non-experts could apply to their own data, without the need for specialized hardware.

Stacked classifiers have been shown to perform well when additional, domain-specific features are derived from the prediction of the classifier at each level in the stack. For example, in the Auto-context model, new features are generated by sampling predictions over a contextual grid, and as a result the stacked classifier is able to learn stereotypical class layouts [26]. This approach has been applied to numerous different tasks, including facade segmentation [11], and bio-medical image segmentation [13, 18]. Here we adapt this idea to address the challenge of splitting touching nuclei by introducing “circularity features” that leverage prior knowledge of nucleus shape to better identify their boundaries.

There are numerous software packages for creating pipelines with segmentation and detection, such as FIJI [21], Cell Profiler [5], ICY [8] and many more. We opted to construct assistive bots in Matlab based on (i) our pre-existing code base in Matlab for puncta detection, (ii) rapid development and quantitative evaluation of circularity features in the stacked Random Forest model, and (iii) ease of developing custom bots for every step in the workflow, such as annotation and parsing of annotations into label maps.

## 3. Workflow and Algorithms

Figure 1 illustrates a typical workflow in SpotsInNucleiBot (SNB). In essence there are four tasks: (1) load stack, (2) set up segmentation parameters, (3) set up spot detection parameters, (4) score folder.

We provide two algorithms for Task (2): Simple Threshold (which is described in Subsection 3.1), and Machine Learning (Subsection 3.2). We also provide two algorithms for Task (3): LoG (as described in Subsection 3.3) and Advanced LoG (Subsection 3.4).

Each choice leads to subsequent mini-apps where parameters can be tested, except for the Machine Learning option, where the user is asked to load a trained model for nuclei segmentation. Such training (and related setting of parameters) are performed in the accompanying NucleiSegmentationBot (NSB).

In summary: for easy nuclei segmentation cases, the user can perform all scoring with SNB. When nuclei segmentation is difficult, the user first trains a nuclei segmentation model via NSB, then loads that model by choosing the Machine Learning option when setting up scoring in SNB.

### 3.1. Basic Nuclei Segmentation

1. Let *I* be the nuclei channel. Set *I*′ = *g_σ_* * *I*, where *g_σ_* is a Gaussian kernel of standard deviation *σ*.
2. Set *I*” = (*I*′ > *t_b_*), i.e., *I*″ is the mask corresponding to pixels in *I*′ with value above *t_b_*. This is the fore-ground (nuclei minus contours) mask.
3. Set *D* as the distance transform of 1 – *I*″.
4. Set *S* = *n*(–*D*), where *n* is the linear mapping to [0,1]. *S* is the raw surface for watershed segmentation, to be refined in the next steps.
5. Set *S*′ = *h_t_h__* (*S*), where *h_t_h__* is the H-minima transform [24] with parameter *t_h_*.
6. Set *S*′(1 – *I*″) = –∞ (here we are using *logical indexing* notation, i.e., setting *S*′(*x*) = –∞ for every pixel *x* where *I*″ = 0). This lets the watershed algorithm know that the background class should be separate from the nuclei object classes.
7. Apply the watershed algorithm on *S*′.

### 3.2. Supervised Nuclei Segmentation

#### 3.2.1 Multi-Layer Classifier

Let {(*I_n_*, *L_n_*): *n* = 1, …, *N*} be a training set, i.e., a set of image/label pairs, where *I_n_*, *L_n_* are 2D tensors, *I* a grayscale image, and *L* a corresponding label map. We use three labels: background, nucleus contour, and nucleus.

The goal is to learn a multi-layer classifier *f* that predicts *L* given *I*. The multi-layer classifier is composed of a sequence of intermediate classifiers *f^i^*, where the superscript *i* indicates the layer index. Each intermediate classifier *f^i^* attempts to predict *L_n_* based on two types of features: *image* features {*F*}^*i*^ – computed directly from *I_n_*, and *probability map* features {*G*}^*i*^ – computed from the predictions {*P*}^*i*–1^ of classifier *f*^*i*–1^. The notation {*X*} indicates a set of distinct features, each of which is referred to as *X*. For example {*F*}^1^ is a set of features computed from an input image, such as derivatives at different directions and scales.

The procedure for training a classifier is as follows:

1. Compute image features {*F_n_*}^1^ from *I_n_*, ∀*n*. Learn a classifier *f*^1^ to predict *L_n_*. The classifier outputs probability maps 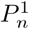.
2. Compute probability map features {*G_n_*}^2^ from 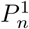, ∀*n*. Learn a classifier *f*^2^ to predict *L_n_*, based on *both* {*F_n_*}^2^ and {*G_n_*}^2^.
3. Repeat step 2 recursively. The number of executions of step 2, plus one, is referred to as the number of the *layers* of the stacked classifier.
4. Convert the final probability map {*P_n_*}^*î*^ to *objects* (see 3.2.3 for details). Here *î* is the index of the last layer.

After tests with a number of different image features, and considering the trade-off between classifier accuracy and computational cost, we settled on features based on image derivatives, up to second order, in different scales. Using 5 different scales (spaced exponentially), this corresponds to 40 features (for each scale there are 8 features: *I*, *∂_x_I*, *∂_y_I*, *∂_xx_I*, *∂_xy_I*, *∂_yy_I*, and the two eigenvalues of the Hessian matrix of *I*). Notice that some of these features, such as the eigenvalues and second derivatives, capture some geometry of the nuclei shapes. For example, at large scales, nuclei are seen as local maxima of the second derivatives. These are, however, different from circularity features (see 3.2.2), which are meso-scale shape features, with the scale set by the size of the objects.

We compute three types of features from probability maps: edge likelihoods, offset features, and circularity features. Edge likelihoods^4^ are computed from the *fore*-*ground* probability maps, meaning the probability maps corresponding to nuclei (interior, not contour). Edges are computed via wavelets (according to the maximum magnitude response of a bank of wavelets with fixed magnitude and different orientations equally spaced in [0, 2*π*). Figure 3 (c) shows an example.

Offset features are computed as in [26]. For a set of offsets *d* ∈ {*d*_1_, …, *d_M_*} and angles *α* = {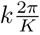: *k* = 0, …, *K* – 1}, we translate 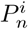 by *d* · (cos *α*, sin *α*). We used a small number (3) of offsets *d* up to the estimated radius of a small nucleus, and set *K* = 8. This adds 72 features to the model.

Finally, we add circularity features, as defined in the next subsection.

#### 3.2.2 Circularity Features

Let 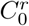 (*x*, *y*) be the likelihood that there is a circle of radius r centered at coordinates (*x*, *y*) in the image. 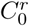 can be computed via the circular Hough Transform, or any of its variations (we use [6]).

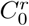 is not a good feature for nearly (including perfectly) circular objects because it is only prominent at the center of such objects – ideally we want a feature that labels all pixels in a circle, or its boundary, not just the center. But from 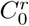 we can infer likelihoods of *boundaries* (which we call *circumference* likelihoods), by accumulating drawings of circumferences of radii *r* centered at (*x*, *y*) with strength (pixel value) 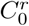. Similarly, we can infer the likelihoods of interiors (*circle* likelihoods), by accumulating drawings of circles.

To improve computation time, one can choose to use only center likelihoods 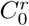(*x*, *y*) above a certain threshold, and only draw a “dense enough” subset of interior (circle) pixels for each chosen center likelihood, and use blurring to “fill in” the gaps. This choice depends on computation time and on how well the feature reconstructs the circle/circumference. In addition, we can add circles and circumferences for multiple center likelihoods (of different radii) in the same accumulator space. The entire algorithm is shown in Figure 2, and some examples are shown in Figure 3.

**Figure 2.**
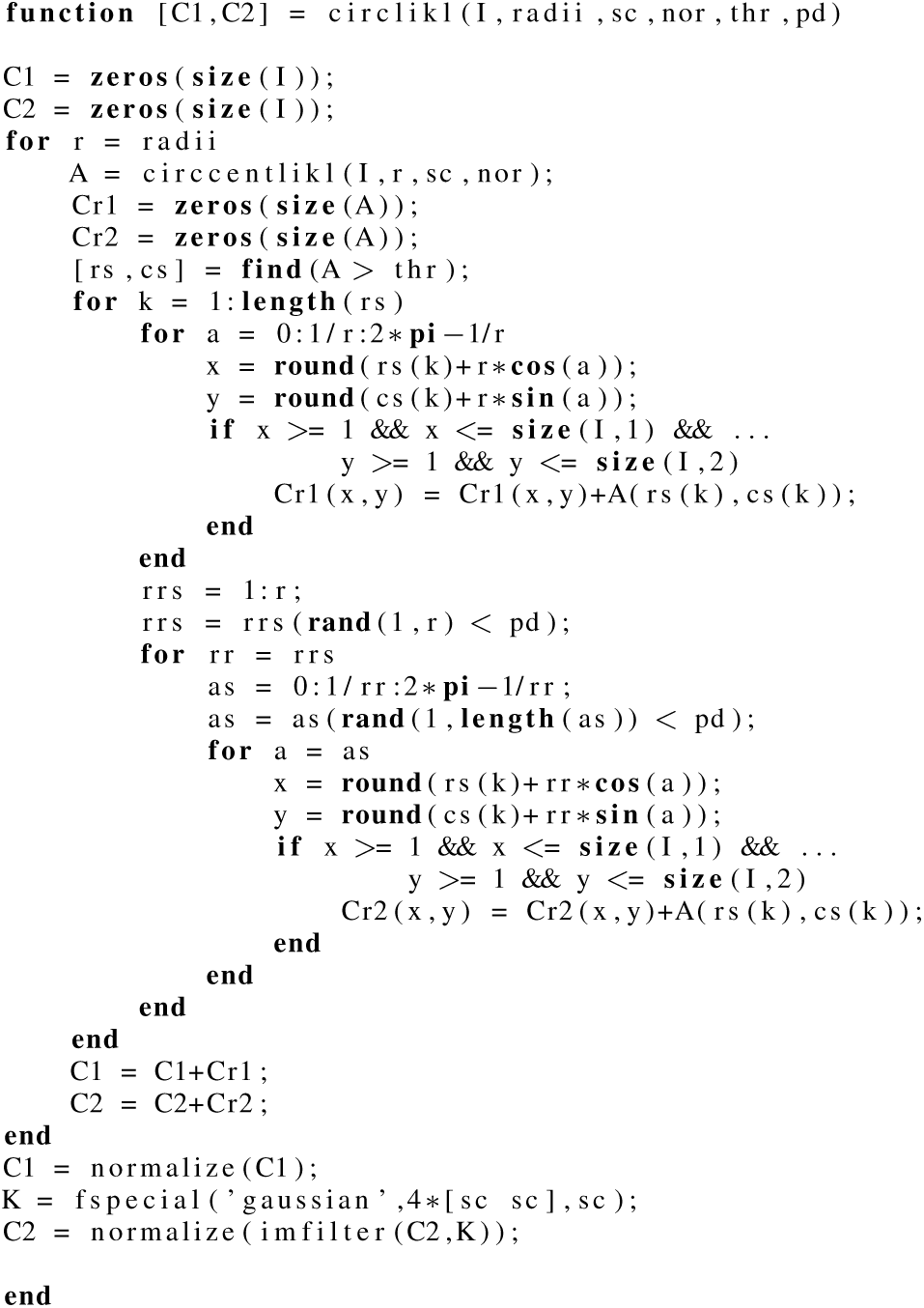
Circularity features algorithm in Matlab. **Inputs.** `I`: image; `radii`: list of radii; `sc`: scale of wavelets used in `circcentlikl` (line 6) and for smoothing circle accumulator space (line 39); `nor`: number of orientations used by `circcentlikl`; `thr`: threshold for contributing with circumference/circle; `pd`: pixel density for drawing circle. **Outputs.** `C1`: circumference likelihood; `C2`: circle likelihood. **Variation.** In the applications we actually replace line 34 with `C1` = `C1`+`Cr1`–`A`: this in some case helps reducing clutter at the center of circles. **Dependencies.** `circcentlikl`, as the name suggests, computes circle center likelihoods (see [6]); `normalize` sets the image into range [0,1].

**Figure 3.**
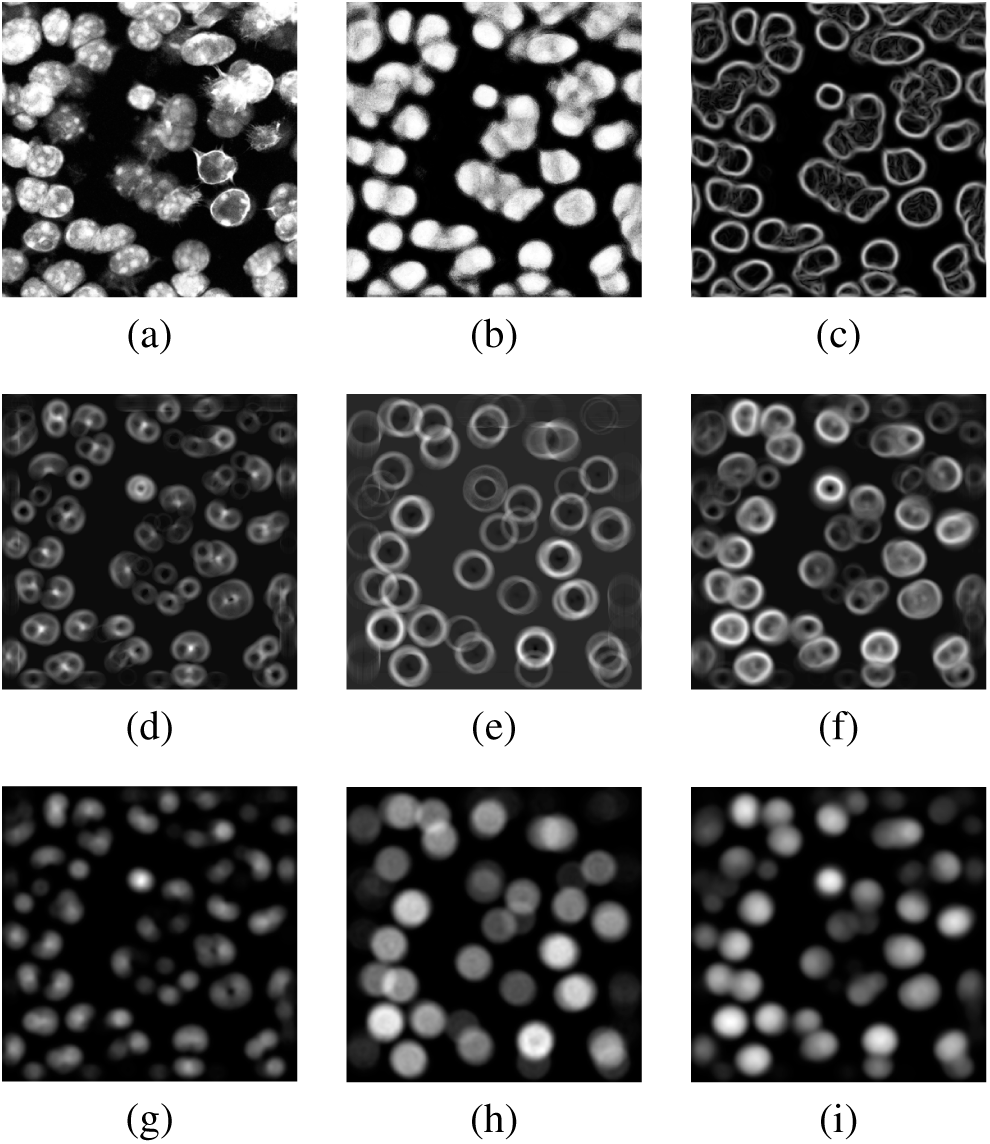
Probability map features. (a) nuclei image; (b) fore-ground (nuclei minus contours) probability map; (c) edge like-lihood; (d,e,f) circumference like-lihoods for *r* = 10, *r* = 20, *r* = 10, 11, …, 20; (g,h,i) circle likelihoods for the respective radii. Notice how circumference features do a better job at highlighting the boundary between touching nuclei than edge features.

In the experiments, we used a total of 3 pairs (pair = circle+circumference) of circularity features for 3 radii around the estimated radius of a nucleus.

#### 3.2.3 From Probability Maps to Objects

We now give more details on step 4 of the pipeline described in Subsection 3.2.1.

For simplicity, let’s call *P* the final probability map of step 3 in that pipeline. *P* is a 3D tensor, [*P*_*i*,*j*,*k*_: *i* = 1, …, *m*; *j* = 1, …, *n*; *k* = 1,2,3}, where *m* and *n* are the height and width of the image, respectively. Lets call *P_k_* the k-th slice of *P* along the 3rd dimension: *P_k_* = {*P*_*i*,*j*,*k*_∀*i*, *j*}. Thus, *P*_1_, *P*_2_, and *P*_3_ are probability maps for the background, nucleus contour, and nucleus body classes, respectively.

Segmented objects are generated from *P* as follows:

1. Set 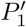:= *σ*(*P*_1_), where *σ*(·) = *g_σ_* * (·), is convolution with a Gaussian kernel with standard deviation 2 pixels.
2. Set 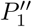:= (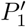 > *t_b_*), i.e., 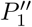 is the mask corresponding to pixels in 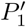 with value above *t_b_*. This is the background mask.
3. Set 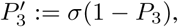 where *σ*(·) is as above.
4. Set 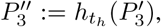 where *h_t_h__* is the H-minima transform [24] with parameter *t_h_*.
5. Set 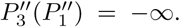 This lets the watershed algorithm know that the background class should be separate from the nuclei object classes.
6. Apply the watershed algorithm on 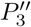.

Notice that *P*_2_, the contour probability map, is not used during post-processing. It still has an important role, however: we found that the Random Forest classifier produces better foreground (nuclei minus contours) maps when trained with three classes (background, nuclei contour, nuclei interior) than when trained with only two (background and nuclei).

### 3.3. Basic Puncta Detection

When the signal-to-noise ratio is high, a simple algorithm based on the Laplacian of Gaussian (LoG) filter performs quite well. It works as follows.

First we filter the puncta channel with a LoG kernel of appropriate sigma (the SNB allows the user to estimate sigma by simply drawing a box around a few puncta). Then the intensities of the local maxima of the filtered image are divided in two groups: those that intersect the nuclei mask, and those that don’t. Robust mean *m* and standard deviation *s* of the background puncta are computed. Finally, all local maxima with intensities above *m* + *ts*, where *t* is a user defined threshold, are considered puncta.

### 3.4. Advanced Puncta Detection

For low SNR cases we include a simplified version of the method described in [1], where the amplitude and background are regressed at each local maximum by estimating the amplitude of a fitting Gaussian centered on that location. (For computational speed we omit the second step in which all parameters of such fitting Gaussian are estimated.)

## 4. Nuclei Segmentation Experiments

In this section we describe experiments conducted to evaluate the performance of the stacked Random Forest method on segmenting nuclei. Other algorithms were not evaluated either because they are standard for simple cases (nuclei detection via thresholding and watershed, and LoG spot detection) or have been published/evaluated elsewhere [1].

The multi-layer Random Forest with circularity features (RF+) was compared against a similar version without circularity features (RF), as well as U-Net, a state-of-the-art Deep Learning architecture for semantic segmentation [20].

From a labeled set of 23 images containing a total of about 2000 nuclei from slices of mouse cerebral cortex, we obtained 93 non-overlapping patches (each with 360×360 pixels), which were split into training (60 images), validation (10 images) and test (23 images). This database is publicly available^5^.

### 4.1. Evaluation

Nucleus segmentation is an instance-segmentation task: not only do we want to classify each pixel as belonging to a nucleus or not, we also want to aggregate nucleus pixels into individual instances. Critically, touching nuclei should be identified as separate instances. In these types of problems, an evaluation metric based on *intersection over union* is best suited to evaluate performance.

Here we adapt what in [23] is defined as the Weighted Coverage Score, normalizing it by the maximum weighted coverage score per image, so that the maximum is 1. We call this measure the *Normalized Coverage Score*. It is defined as follows:

Let 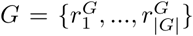 be a set of ground truth regions and 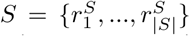 be a set of proposed regions for a given image. For a given pair of regions *r_j_* and *r_k_*, the overlap between them is defined using the intersection over union score: *O*(*r_j_*, *r_k_*) = (*r_j_* ∩ *r_k_*)/(*r_j_* ∪ *r_k_*). The *weighted* coverage score, as defined in [23], is given by

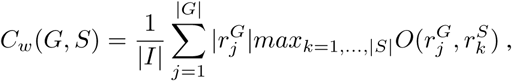

where |*I*| is the total number of pixels in the image and 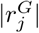 is the number of pixels in the ground truth region 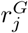. The *normalized* coverage score is defined here as

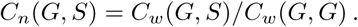

Simple algebra shows this is equivalent to

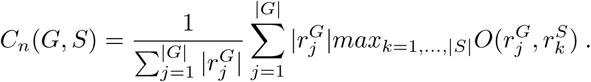

### 4.2. Results

For each combination of parameters 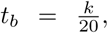 *k* = 1, …, 9, and 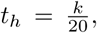 *k* = 1, …, 4 in the watershed post-processing phase (Subsection 3.2.3), we computed coverage scores for three classifiers: 3-layer Random Forests with (RF+) and without (RF) circularity features, and a U-Net with 3 downsampling (and upsampling) layers. We implemented our own modified version of the U-Net so that the output image is the same size as the input image. The outputs (and ground truth) are filtered to remove: (i) objects that touch the border of the image, (ii) objects that are smaller than a circle of radius half the smallest expected circle or larger than a circle of radius twice the largest expected circle (such filtering alters the labels only minimally, excluding small artifacts that appear at the intersection of neighboring nuclei due to imprecise annotation), (iii) objects with more than 10% of their area intersecting a region marked to be ignored.

Figure 4 shows coverage scores for fixed *t_h_* and varying *t_b_*, with background sampling restricted to regions near nuclei boundaries. Figure 5 shows a comparison of the outputs of the three models.

**Figure 4.**
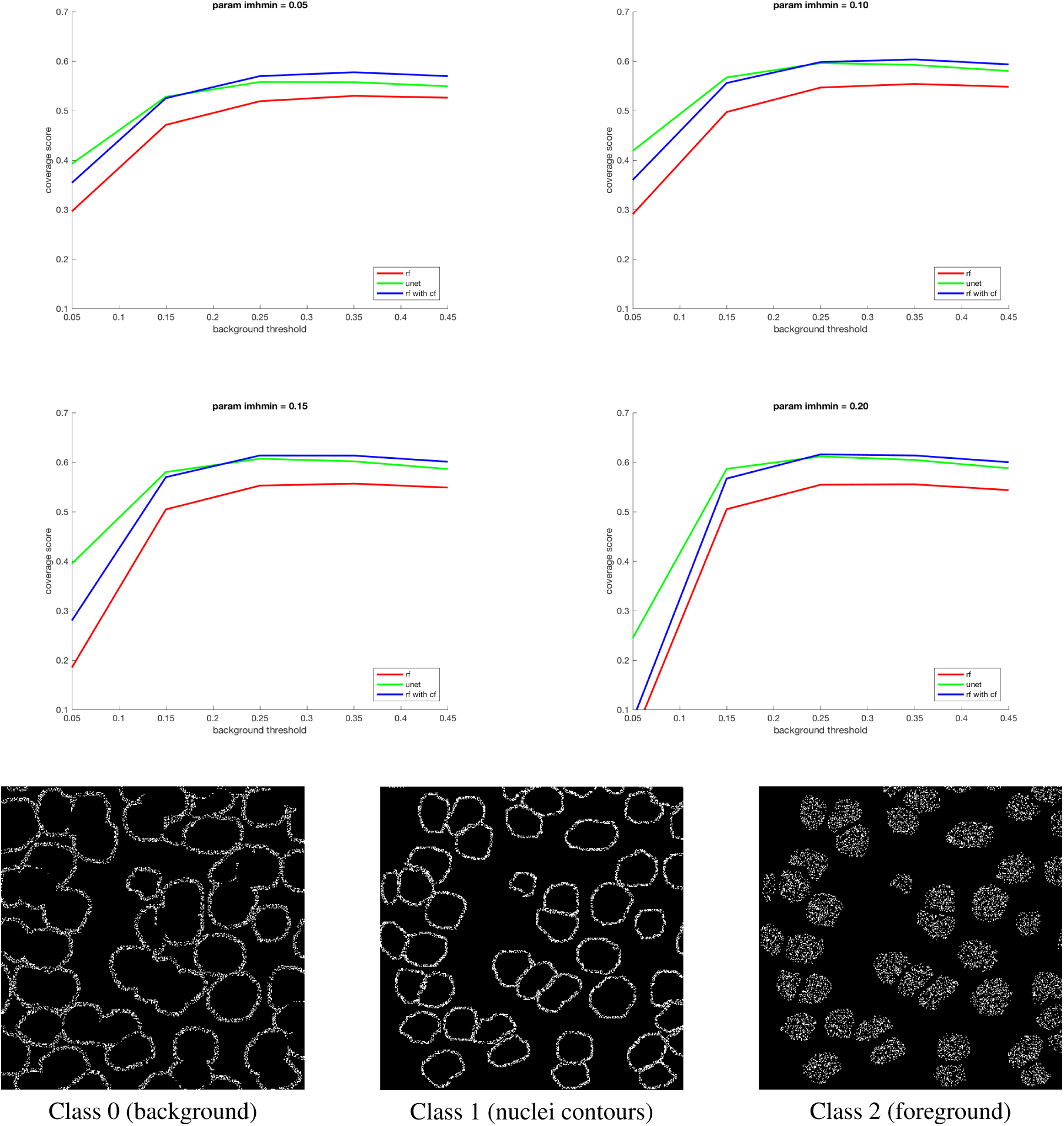
Normalized coverage scores on the test set for models trained with a balanced number of samples per class, and background samples taken near the boundary of nuclei (as exemplified in the 3 bottom images).

**Figure 5.**
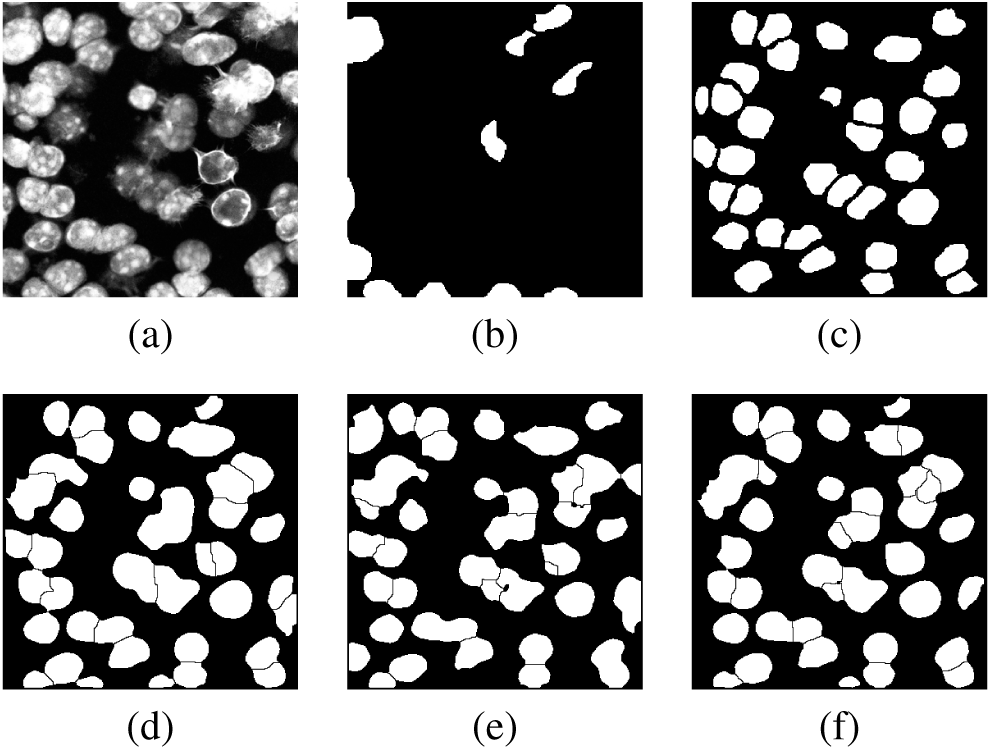
Example of segmentation outputs using different models, trained with class balancing, and background class sampled near nuclei boundaries, with post-processing parameters *t*b** = 1/4 and *t_h_* = 1/5. (a) nuclei image; (b) ignore labels; (c) nuclei labels; (d) output of Random Forest without circularity features; (e) output of U-Net; (f) output of Random Forest with circularity features.

We found that 3-layer Random Forests reached the optimal trade-off between evaluation speed and accuracy on our data, and the same held true for 3-layer U-Nets. For brevity, we do not report results on other architectures.

### 4.3. Applications

SpotsInNucleiBot is designed to work on any multichannel images containing, in one channel, roughly circular nuclei, and in the remaining channels, spot-like objects (spots, point-sources, puncta) that can be approximated by a 2D Gaussian of small standard deviation. To demonstrate this flexibility, Figure 6 illustrates our approach applied to two distinct bio-image analysis problems: RNA detection in slices of mouse visual cortex, and protein detection in human fibroblasts.

**Figure 6.**
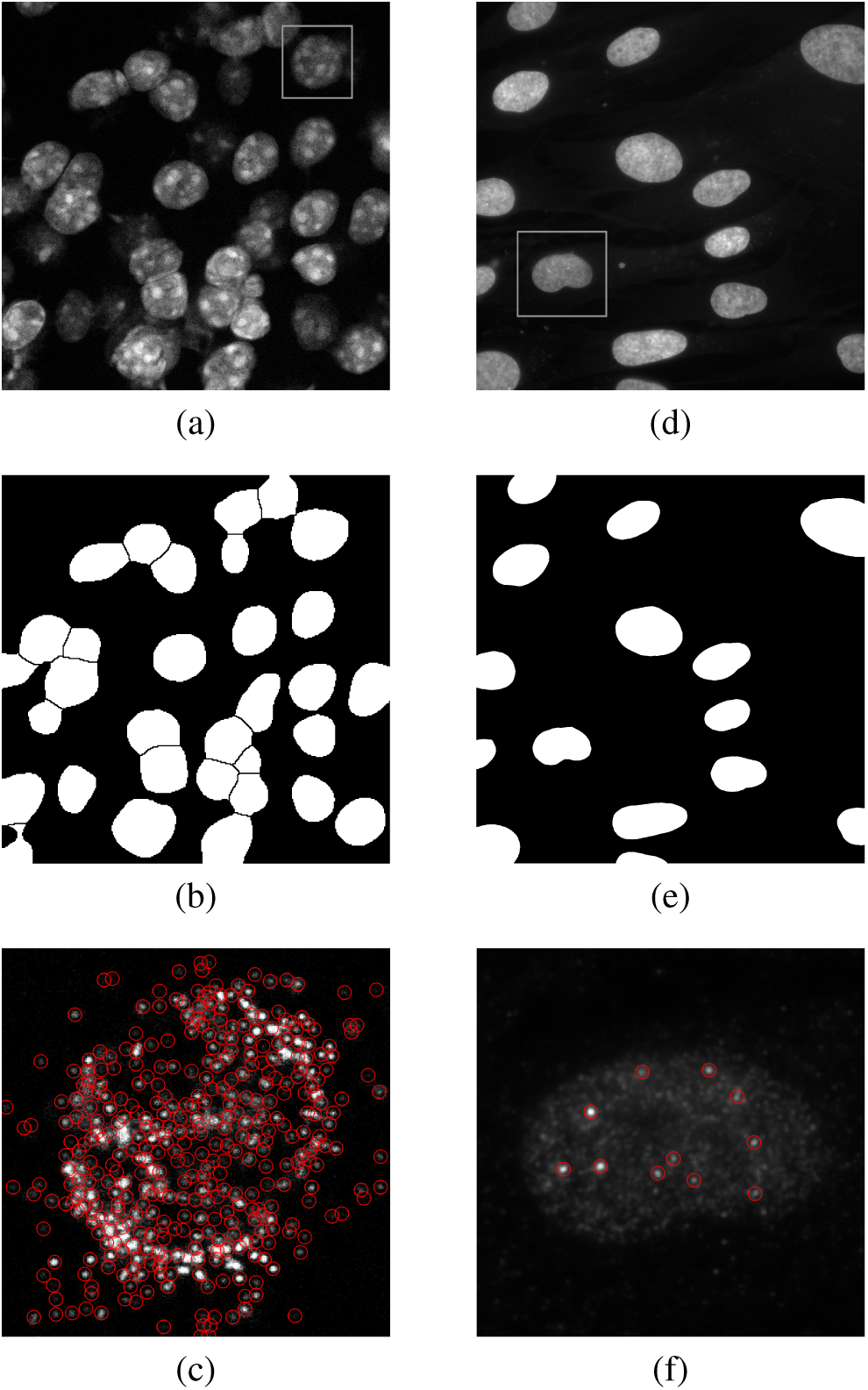
Applications to diverse samples and probed substrates. Left column: a slice of mouse visual cortex (a) and the output nuclear segmentation (b) and puncta detection of fluorescently labeled *Egrl* RNA (c) for selected ROI shown in (a). Right column: Human fibroblasts (d) and the output segmentation (e) and protein puncta detection with a fluorescently labeled antibody for ATRX protein (f) for selected ROI shown in (d). Nuclei segmentation in (b) is performed via Machine Learning, and in (e) via Simple Threshold. Spot detection in (c) uses the Advanced LoG option, while simple LoG was used in (f).

## 5. Conclusion

The task of segmenting nuclei and detecting associated fluorescently-labeled probes for DNA, RNA, or protein from light micrographs arises frequently in bio-image analysis, and is essential for the rapidly growing field of image-based transcriptomics. We present a user-assistive approach to address this problem, based on the use of “bots”. Specifically, we provide bots that assist in the process of segmenting nuclei, estimating model parameters for puncta detection, detecting puncta, and associating puncta to their corresponding nuclei.

Accurate segmentation of nuclei is particularly challenging due to the difficulty of separating tight clusters of nuclei commonly found in complex tissues. For these more difficult segmentation scenarios, we provide a new method based on stacked Random Forests. We introduce *circularity features* to improve separation of touching objects, based on an implicit shape model. This approach achieves comparable performance to the state-of-the-art Deep Learning approach [20], without the requirement for specialized hardware, empowering non-experts to train and apply this Machine Learning model on their own data.

The use of bots provides a flexible framework for incorporating new methods as they are developed by the large academic community of computer vision and bio-image analysis experts programming in Matlab. Bots for detection of other biologically relevant substrates could be easily integrated into this pipeline, and new pipelines can be quickly assembled using our segmentation and detection bots.

## 6. Acknowledgements

We would like to acknowledge the Harvard Medical School Tools and Technology fund, as well as the William F. Milton fund, for supporting this research. We also thank Joseph Cabral and David Knipe, Department of Microbiology and Immunobiology, Harvard Medical School, for providing the micrographs in Figures 6 (d,f), prepared as a part of research supported by NIH grant AI106934.

---

**Supplementary Figure 1.**
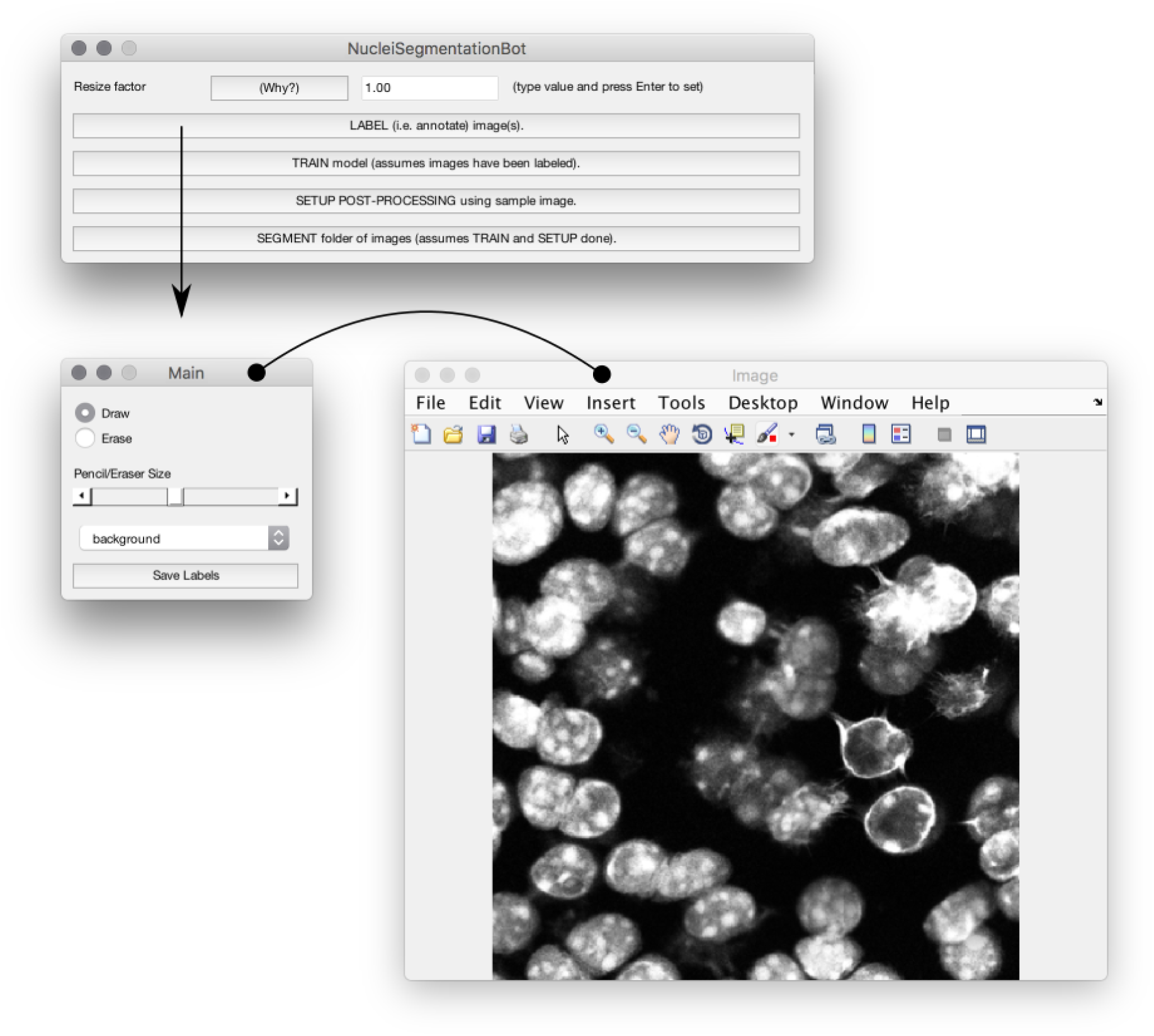
NucleiSegmentationBot sample workflow. In this particular section of the bot the user can annotate background-class pixels. Notice that the Main and Image windows belong to the same mini-app for annotation. Note: not shown here are intermediate steps/windows related to navigating to the image to be opened for annotation (after the button LABEL is clicked).

**Supplementary Figure 2.**
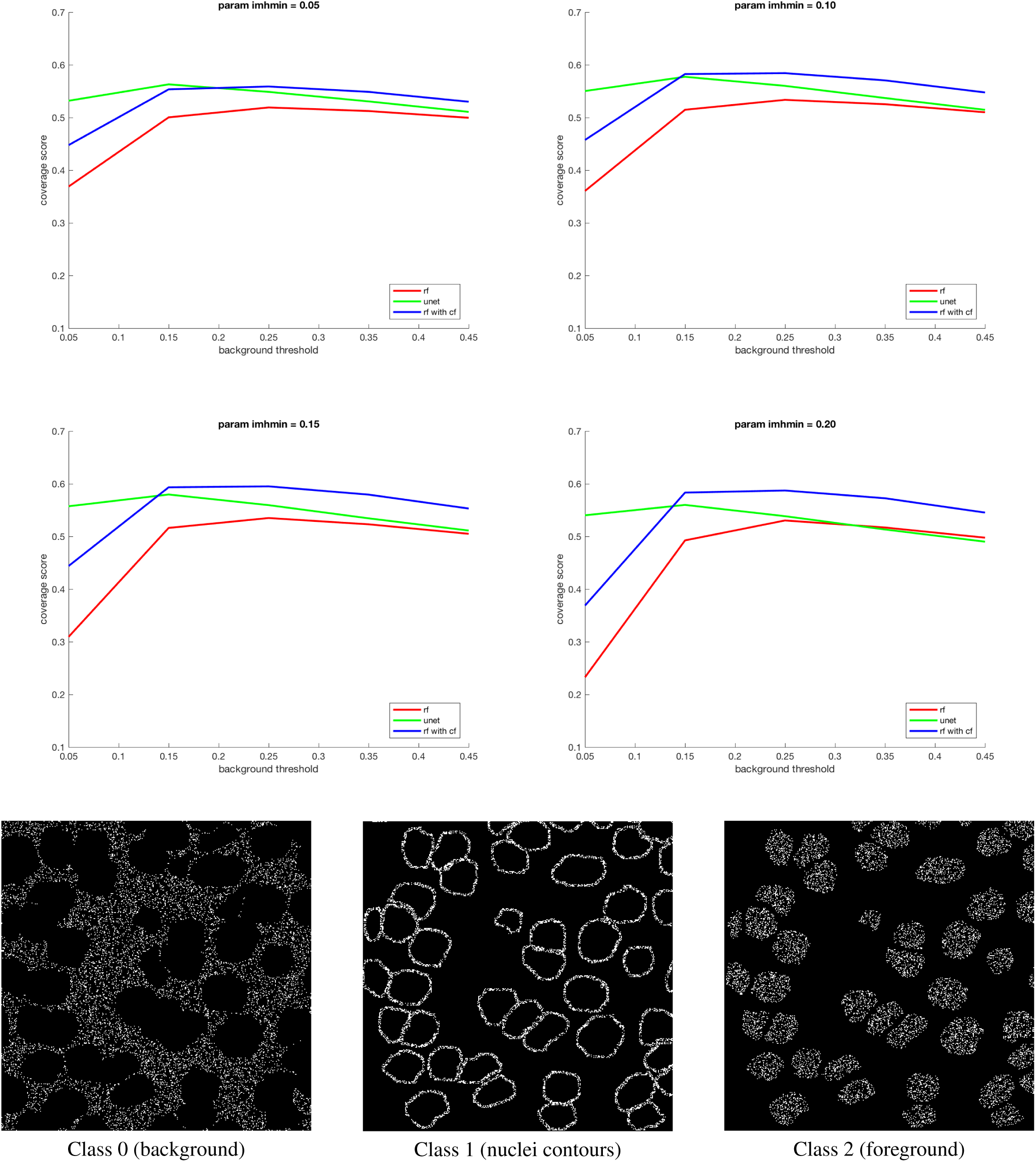
Normalized coverage scores on the test set for models trained with a balanced number of samples per class, and samples taken without regard for location in the image (as exemplified in the 3 bottom images).

## Footnotes

1 Scoring refers to counting the number of puncta, potentially in multiple different channels, associated with each nucleus. Puncta refers to point-like objects, which can be diffraction limited or not. When diffraction limited, they are also sometimes called “point sources”.

2 Code, sample datasets, and video tutorials, are available at https://hms-idac.github.io/MatBots/

3 https://hms-idac.github.io/MLRFSwCF/

4 The term “likelihood” is sometimes used instead of “feature.” A likelihood is simply a feature with some geometric meaning. For example, edge likelihood is a feature where higher values represent higher likelihood that there’s an edge at the corresponding point in the image. The edge likelihood is a combination of features given by edge filters in all directions.

5 https://hms-idac.github.io/MLRFSwCF/

